# Non-selective inhibition of the motor system following unexpected and expected infrequent events

**DOI:** 10.1101/2020.03.25.008789

**Authors:** Carly Iacullo, Darcy A. Diesburg, Jan R. Wessel

## Abstract

Motor inhibition is a key control mechanism that allows humans to rapidly adapt their actions in response to environmental events. One of the hallmark signatures of rapidly exerted, reactive motor inhibition is the non-selective suppression of cortico-spinal excitability (CSE): unexpected sensory stimuli lead to a suppression of CSE across the entire motor system, even in muscles that are inactive. Theories suggest that this reflects a fast, automatic, and broad engagement of inhibitory control, which facilitates behavioral adaptations to unexpected changes in the sensory environment. However, it is an open question whether such non-selective CSE suppression is truly due to the unexpected nature of the sensory event, or whether it is sufficient for an event to be merely infrequent (but not unexpected). Here, we report data from two experiments in which human subjects experienced both unexpected and expected infrequent events during a simple reaction time task while CSE was measured from a task-unrelated muscle. We found that expected infrequent events can indeed produce non-selective CSE suppression – but only when they occur during movement initiation. In contrast, unexpected infrequent events produce non-selective CSE suppression even in the absence of movement initiation. Moreover, CSE suppression due to unexpected events occurs at shorter latencies compared to expected infrequent events. These findings demonstrate that unexpectedness and stimulus infrequency have qualitatively different suppressive effects on the motor system. They also have key implications for studies that seek to disentangle neural and psychological processes related to motor inhibition and stimulus detection.

## Introduction

Motor inhibition is a core component of controlled and flexible human behavior. The rapid interruption of active motor representations allows humans to momentarily cancel ongoing movements and movement plans, which in turn allows them to reevaluate whether those movements are still appropriate when environmental circumstances suddenly change. In the laboratory, motor inhibition is usually assessed in tasks like the stop-signal task (Logan, Cowan, and Davis, 1984), where it allows humans to rapidly stop actions even after their initiation. In such tasks, subjects are explicitly instructed to stop an action following a previously instructed infrequent signal, which follows the response prompt on a minority of trials (Verbruggen et al., 2019). Because subjects in tasks like the stop-signal task expect that these infrequent stop-signals will occur on a subset of trials, successful action-stopping in such tasks results from the implementation of both proactive and reactive inhibitory control mechanisms (Aron, 2011, Kenemans, 2015). Proactive inhibition denotes the anticipatory implementation of control processes during the expectation of a stop-signal, while reactive inhibition denotes the cascade of processes that is triggered by the stop-signal itself (Verbruggen et al., 2009; Chikazoe et al., 2009; Jaffard et al., 2008).

Within the stop-signal task, the signals that instruct participants to cancel an action are infrequent events. However, since stop-signals are explicitly part of the task instruction, their occurrence is also *expected*. Notably, however, in recent years, work on tasks that involve *unexpected* sensory events (e.g., the novelty-oddball paradigm or the cross-modal oddball task; Courchesne et al., 1975, Parmentier et al., 2008) has shown that such events automatically induce motor inhibition, even when there is no instruction to ever stop an action. In other words, unexpected sensory events induce a reflexive engagement of motor inhibition, and they can do so even in the absence of proactive control (i.e., when the task does not involve an instruction to exert inhibitory control; Wessel, 2018).

This automatic recruitment of reactive motor inhibition after unexpected events is evident on many levels of observation, including behavior, brain activity, and physiological changes of the motor system (cf. Wessel & Aron, 2017, for a review). In behavior, this engagement of motor inhibition is suggested by the fact that unexpected events presented during forced-choice reaction time tasks lead to a slowing of the prompted motor responses (Dawson et al., 1982, Ljungberg et al., 2012). Concomitantly, in the brain, unexpected events activate some of the same cortical and subcortical circuitry that is involved in stopping actions in tasks like the stop-signal task (Bockova et al., 2011; Wessel et al., 2016; Fife et al., 2017).

However, the inhibitory effects that unexpected events exert on the motor system are perhaps most evident from physiological measurements of cortico-spinal excitability (CSE). CSE can be non-invasively probed using transcranial magnetic stimulation (TMS) and electromyography (Barker et al., 1985; Rothwell et al., 1999; Bestmann & Krakauer, 2015). By applying single-pulses of TMS to the contralateral motor cortex representation of a specific muscle, a motor evoked potential is produced in the electromyogram of that muscle. The amplitude of this motor evoked potential provides a proxy for the net-CSE of the underlying corticomotor tract. In tasks like the stop-signal task, CSE of the muscles involved in the action is suppressed when a stop-signal occurs (Coxon et al., 2006, 2007). In addition, several studies have shown that this suppression of the motor system extends even beyond the muscle group that is targeted for stopping (Badry et al., 2009; Cai, Oldenkamp, and Aron, 2011; Majid et al., 2013; Wessel et al., 2013, 2016). Subsequent studies have found that the proactive-reactive control balance is a key factor in determining this non-selective property of motor inhibition: the more proactive control is exerted, the more selectively it can be applied. In turn, the more stopping relies on reactive mechanisms, the greater the non-selective suppression of CSE (Greenhouse, Oldenkamp, and Aron, 2012, Duque et al., 2017). In other words, non-selective CSE suppression is a hallmark signature of the reactive implementation of motor inhibition. Consequently, in line with the proposal that unexpected sensory events lead to an automatic recruitment of the brain’s reactive inhibition circuity even when stopping is not explicitly required (i.e., in the absence of proactive control), such events do indeed also produce non-selective suppression of CSE (Wessel & Aron, 2013). In that particular study, subjects performed a verbal reaction time task, in which unexpected sounds were infrequently presented prior to the imperative stimulus. This led to CSE suppression at a task-unrelated hand muscle, specifically at 150ms following sound onset. The same is true when a task is performed with the legs and CSE is measured at the hand (Dutra et al., 2018).

Such studies of unexpected sensory events (see also Novembre et al., 2018, 2019) have led us to propose that unexpected events automatically activate the same reactive inhibitory control systems that are recruited when actions have to be stopped actively in tasks like the stop-signal task. Specifically, we propose that the purpose of this automatically engaged inhibitory control effort is to rapidly interrupt ongoing behavior, which purchases time for the cognitive system to resolve the surprise produced by the unexpected event. This additional processing time can be used to evaluate whether ongoing motor plans are still appropriate in light of the sudden unexpected change in environmental regularity (Wessel & Aron, 2017).

However, there is a notable alternative to this surprise-inhibition theory. Specifically, while the two types of psychological events that are known to result in non-selective CSE suppression (stop-signals and unexpected events) differ in the degree to which they produce surprise (stop-signals are expected, unexpected events are not), they also have a notable commonality: they are both *infrequent* events within the context of their respective tasks. Stop-signals typically occur in around 25-33% of trials in the stop-signal task (Verbruggen et al., 2019; though see Dykstra et al., 2020 for an exception). Similarly, in typical studies of unexpected events, only about 10-20% of trials involve an unexpected event. Therefore, it is possible that the infrequency of a stimulus alone can account for the presence of non-selective CSE suppression after both stop-signals and unexpected events. If that is the case, surprise itself is not necessary to explain the presence of non-selective CSE suppression, and may in fact not uniquely engage motor inhibition at all. Indeed, while surprise and infrequency are often confounded, they are meaningfully different cognitive constructs. For example, infrequent events can be entirely expected (hearing a fire alarm during an announced drill), or entirely unexpected (hearing the same fire alarm on a regular day), with fundamentally different cognitive and behavioral implications.

Therefore, the goal of the current study was to investigate whether *expected* infrequent events can produce reactive motor inhibition, as indexed by non-selective CSE suppression.

Notably, this question is not just relevant to test the proposed link between surprise and motor inhibition. Indeed, if expected infrequent events can – by themselves – recruit reactive motor inhibition, this would be highly relevant for the study of motor inhibition in the stop-signal task. In fact, one of the most controversial questions in the recent stop-signal literature is which exact neural or psychological processes following stop-signals are related to the attentional detection of the infrequent stop-signal itself, and which are related to the actual implementation of motor inhibition (Verbruggen et al., 2010; Hampshire et al., 2010; Matzke et al., 2013). To address this question, many studies have utilized control tasks whose stimulus layout matches the stop-signal task (i.e., a go-signal is followed by an infrequent second signal) but with an instruction that does not involve outright action stopping (e.g., to press a second button after the original go-response or to ignore the second signal entirely, Hampshire et al., 2010; Dodds et al., 2011; Chatham et al., 2012; Erika-Florence et al., 2014; Waller et al., 2019). If such expected infrequent stimuli presented outside of a stop-signal task produced the same type of reactive, non-selective motor inhibition that is found after unexpected infrequent stimuli, it would invalidate the assumption that a contrast between a stop-signal and an infrequent-signal control task would isolate the inhibitory process that is found in the stop-signal task.

Therefore, in sum, we here aimed to explicitly test whether *expected* infrequent events produce the same type of non-selective suppression of the motor system that is found after *unexpected* infrequent events. We tested this possibility using tasks that presented such infrequent events both *before* and *during* action initiation.

## Methods

### Participants

In Experiment 1, participants were twenty young, healthy adults (17 female, mean age 18.65, SD: .9). In Experiment 2, participants were twenty-one young, healthy adults (all right-handed, 14 female, mean age: 20.76, SD: 4.2). All participants were recruited via a University of Iowa research-dedicated email list or via the University of Iowa Department of Psychological Brain and Sciences’ online recruitment tool and compensated in correspondence to their recruitment means, either by an hourly rate of $15 or by receiving course credit. The participants were all screened using a safety questionnaire (Rossi et al., 2011) to ensure it was safe for them to undergo TMS. Experimental procedures were approved by the University of Iowa Institutional Review Board (#201711750).

### Experimental task

The stimuli for the behavioral paradigms for both Experiment 1 and Experiment 2 were presented using Psychtoolbox (Brainard et al., 1997) and MATLAB 2015b (TheMathWorks, Natick, MA) on a Linux desktop computer running Ubuntu. In Experiment 1, participants responded to the stimuli on the screen using their feet by pushing Kinesis Savant Elite 2 foot pedals (left or right; see ***Figure 1*** for visualization of task setup). At the beginning of every trial, a black fixation cross was displayed in the center of a gray screen background. After 500ms, a sound stimulus was played for 200ms, which could be of one of the following conditions: STANDARD (frequent), EXPECTED (infrequent), UNEXPECTED (infrequent). The STANDARD and EXPECTED sounds were sine wave tones of either 600 or 800Hz frequency, counterbalanced across participants. The participant was introduced to the STANDARD and EXPECTED sounds in a practice block prior to the recorded experiment. In the practice block, the EXPECTED sound occurred during 20% of trials, with the remainder being STANDARD sounds. In the actual experiment, the EXPECTED sound occurred on 10% of trials. The UNEXPECTED sounds occurred on 10% of trials and were only introduced during the main experiment, without prior instruction. These novel sounds were 90 bird song samples from European starlings (recorded by Jordan A. Comins), which were matched in amplitude envelope and duration to the sine wave tones. After the sound, on each trial, a single pulse of TMS was delivered with a delay of 125, 150, or 175ms (i.e., centered around 150ms, which was the time point at which the CSE suppression after unexpected sounds was observed in Wessel & Aron, 2013). Subjects were instructed that the sound would cue them to the timing of the appearance of the imperative stimulus. The imperative stimulus was a black arrow pointing left or pointing right, and appeared 500ms after the onset of the sound. Participants responded according to the direction of the arrow by pressing the left or right foot pedal (deadline: 1,000ms). If no response was made in time, “Too Slow!” was displayed on screen in red. After an inter-trial interval of 150, 175, 200, 225, or 250ms (during which the fixation cross was displayed), the next trial began. The practice block lasted 30 trials. During the main block, participants completed a total of 810 trials (567 STANDARD, 81 EXPECTED, 81 UNEXPECTED), divided into 9 blocks separated by self-timed breaks.

**Figure 1.**
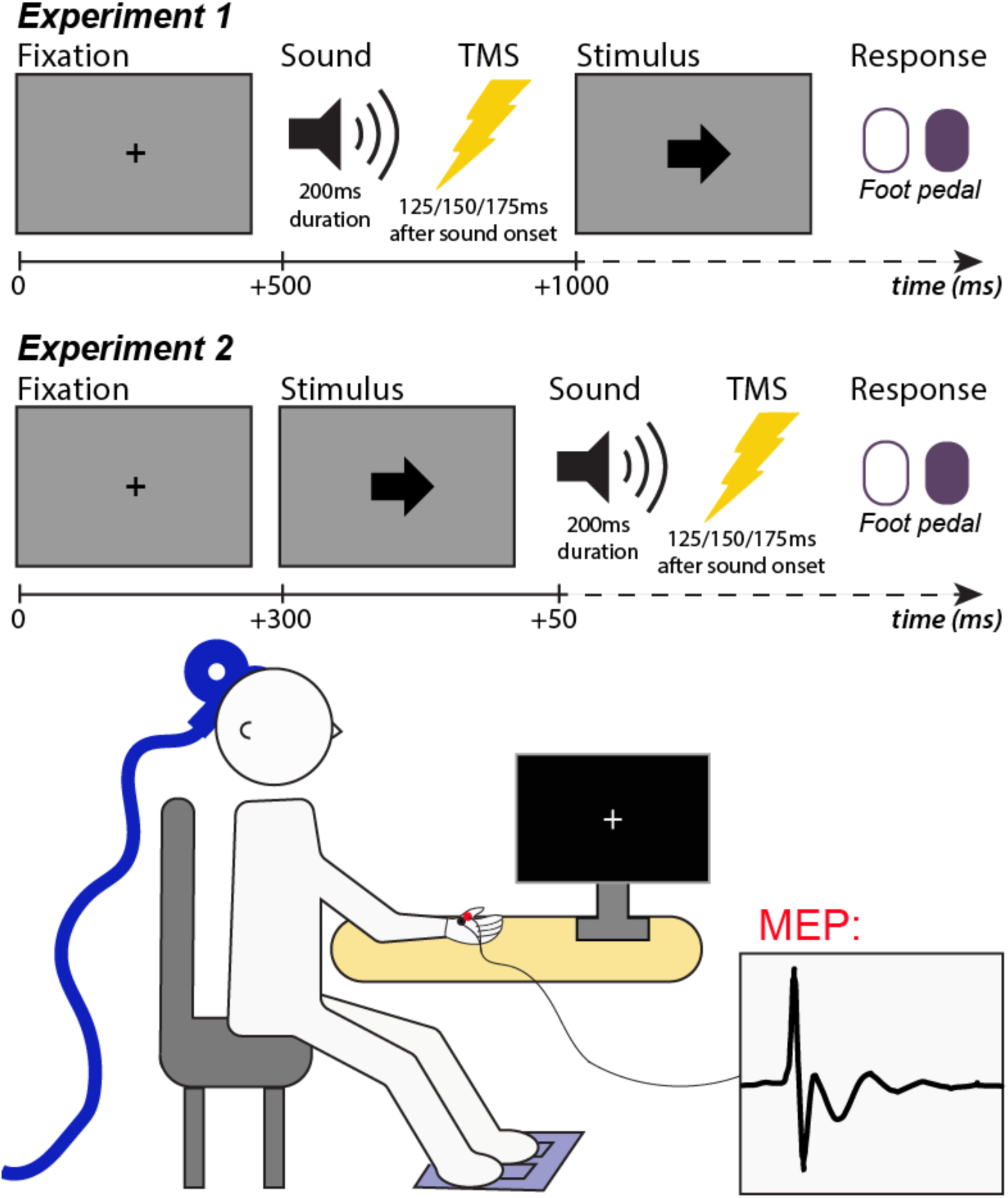
Diagrams of speeded response tasks participants completed in Experiments 1 and 2. In Experiment 1, participants heard the sound before an imperative stimulus (the arrow) was shown. In Experiment 2, participants heard the sound immediately following the arrow. Below the task diagrams is a diagram of the experimental setup: TMS to elicit an MEP is delivered over motor cortex contralateral to the hand muscle with EMG electrodes, while participants respond with the feet.

The task in Experiment 2 was the same as in Experiment 1, except for the order and relative timing of the sound relative to the imperative stimulus. In Experiment 2, the sound played 50ms *after* the onset of the imperative stimulus. Again, TMS stimulation occurred 125, 150, or 175ms after the sound.

All task code, analysis code, and data can be found on the Open Science Framework (OSF) at [link will be added at time of publication].

### TMS protocol

Cortico-spinal excitability (CSE) was measured via motor-evoked potentials elicited by TMS. TMS stimulation was performed with a MagStim 200-2 system (MagStim, Whitland, UK) with a 70-mm figure-of-eight coil. Hotspotting was performed to identify the first dorsal interosseous muscle (FDI) stimulation locus and correct intensity. The coil was first placed 5 cm lateral and 2 cm anterior to the vertex and repositioned to where the largest MEPs were observed consistently. Resting motor threshold (RMT) was then defined as the minimum intensity required to induce MEPs of amplitudes exceeding .1 mV peak to peak in 5 of 10 consecutive probes (Rossini et al., 1994). TMS stimulation intensity was then adjusted to 115% of RMT (Experiment 1: mean intensity: 52.7% of maximum stimulator output; range: 40-68%; Experiment 2: mean intensity: 56.9% of maximum stimulator output; range: 47-67%) for stimulation during the experimental task. In both experiments, TMS pulses occurred with a delay of 125, 150, or 175ms after sound onset (uniform distribution). A passive baseline for MEP normalization was collected by delivery of 10 single TMS pulses at the end of each experimental task block. One baseline pulse was delivered every 3 seconds during baseline collection (the same length as a trial during the active task), and no visual stimuli were shown on screen during this time.

### EMG recordings

An EMG sweep was triggered 90ms before each TMS pulse. EMG was recorded using a bipolar belly-tendon montage over the FDI muscle of the right hand using adhesive electrodes (H124SG, Covidien Ltd., Dublin, Ireland), with a ground electrode placed over distal end of ulna. Electrodes were connected to a Grass P511 amplifier (Grass Products, West Warwick, RI; 1000 Hz sampling rate, filters: 30 Hz high-pass, 1000Hz low-pass, 60Hz notch). The amplified EMG data were sampled via a CED Micro 1401-3 sampler (Cambridge Electronic Design Ltd., Cambridge, UK) and recorded to the disc using CED Signal software (Version 6).

### Motor evoked potential analysis

MEPs were identified from the EMG trace via in-house software developed in MATLAB (TheMathWorks, Natick, MA). Trials were excluded if the root mean square power of the EMG trace 90ms before the TMS pulse exceeded .01 mV or if the MEP amplitude did not exceed .01 mV. MEP amplitude was quantified with a peak-to-peak rationale, measuring the difference between maximum and minimum amplitude within a time period of 10–50 ms after the pulse. Both automated artifact rejection and MEP amplitude quantification were visually checked for accuracy on each individual trial for every data set by a rater who was blind to the specific trial type. We then calculated the mean MEP amplitudes for each condition of interest (*SOUND*: STANDARD, EXPECTED, UNEXPECTED; *TMS TIMING*: 125ms, 150ms, 175ms), and normalized by dividing these amplitudes by the median baseline MEP estimate. After artifact correction, MEP amplitudes were tested for differences using a 3×3 ANOVA with the factors *SOUND* and *TMS TIMING*. For Experiment 1, the mean number of trials per condition were 187 (standard, 125ms), 192 (standard, 150ms), 189 (standard, 175ms), 24 (expected, 125ms), 24 (expected, 150ms), 24 (expected, 175ms), 24 (unexpected, 125ms), 23 (unexpected, 150ms), and 24 (unexpected, 175ms), respectively. For Experiment 2, the mean number of trials per condition were 195 (standard, 125ms), 195 (standard, 150ms), 194 (standard, 175ms), 25 (infrequent expected, 125ms), 25 (infrequent expected, 150ms), 24 (infrequent expected, 175ms), 24 (unexpected novel, 125ms), 23 (unexpected novel, 150ms) and 24 (unexpected novel, 175ms), respectively.

## Results

### Behavior

Mean reaction times and accuracy by condition for both Experiment 1 and 2 can be found in ***Table 1***.

**Table 1.**
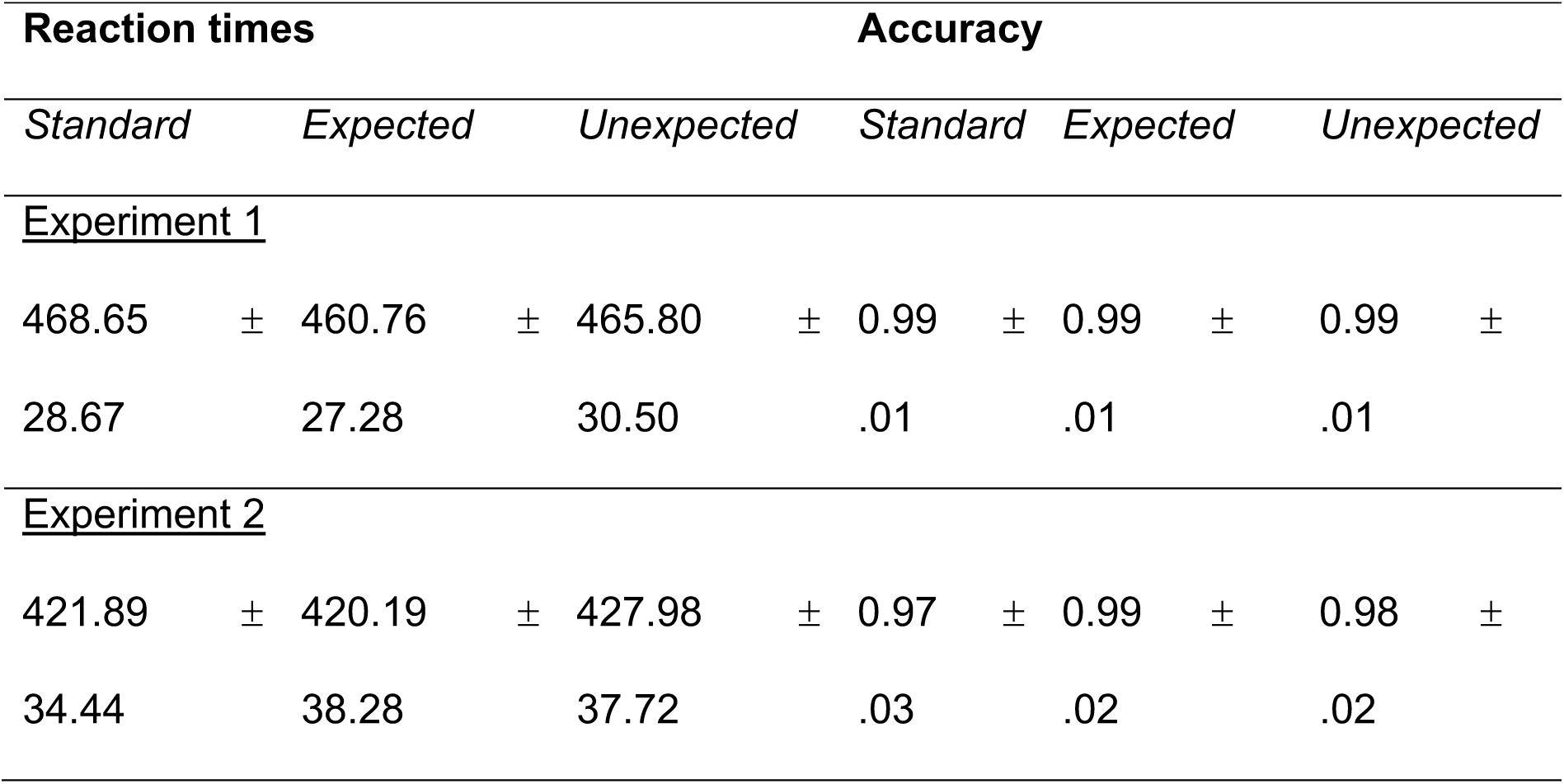
Behavioral results for each sound type from Experiments 1 and 2. Results denote mean +/- standard deviation.

For Experiment 1, we conducted an ANOVA (repeated measures, 1-way/factor) on RT to assess the effects of *SOUND* type. An overall main effect of *SOUND* type was found (*F*(2,19) = 5.03, *p* < .0001, *η*^*2*^ = .96). Pairwise *t*-tests were conducted to evaluate which sound (EXPECTED or UNEXPECTED) resulted in mean RT that differed significantly from RT during the STANDARD trials. Reaction time for UNEXPECTED tone trials was not significantly different from RT during STANDARD trials (*t*(19) = 1.56, *p* = .14, *d* = .09) but RT for EXPECTED tone trials was significantly *faster* than RT during STANDARD trials (*t*(19) = 3.09, *p* = .006, *d* = .28).

In Experiment 2, we presented the sound stimulus following the target arrow to assess the effects of infrequent stimuli on an already-initiated movement. For Experiment 2, we conducted an ANOVA (repeated measures, 1-way/factor) on RT to assess the effects of *SOUND* type. An overall main effect of *SOUND* type was found (*F*(2,19) = 4.66, *p* < .0001, *η*^*2*^ = .96). Pairwise *t*-tests were conducted to evaluate which infrequent sound (EXPECTED or UNEXPECTED) resulted in mean RT that differed significantly from RT during the STANDARD trials. Reaction time for UNEXPECTED trials was significantly slower than RT during STANDARD trials (*t*(20) = -2.77, *p* = .01, *d* = .17), but RT for EXPECTED trials was not significantly different than RT during STANDARD trials (*t*(20) = 0.83, *p* = .42, *d* = .05).

### Cortico-spinal excitability

In Experiment 1 (sound prior to imperative stimulus), no significant main effects of *SOUND* (F(2,19) = 1.57, *p* = .22, *η*^*2*^ = .04) or *TMS TIMING* (F(2,19) = .34, *p* = .72, *η*^*2*^ = .01) were found, and there was no significant interaction of those factors (F(4,19) = .88, *p* = .48, *η*^*2*^ = .05). However, to test whether our previous finding of CSE suppression after UNEXPECTED sounds at the 150ms time point (Wessel & Aron, 2013) replicated, we computed a pairwise *t*-test between MEP amplitudes on STANDARD and UNEXPECTED sounds at the 150ms delay. Indeed, MEPs for UNEXPECTED sounds at 150ms were significantly smaller than MEPs from STANDARD sounds (*t*(19) = 2.05, *p* = .027, *d* = .28, ***Figure 2***). In contrast, the STANDARD vs. EXPECTED comparison showed no significance at any time point (125ms: *t*(19) = 0.72, *p* = .48, *d* = .06; 150ms: *t*(19) = 0.77, *p* = .45, *d* = .09; 175ms: *t*(19) = -0.78, *p* = .44, *d* = .10).

**Figure 2.**
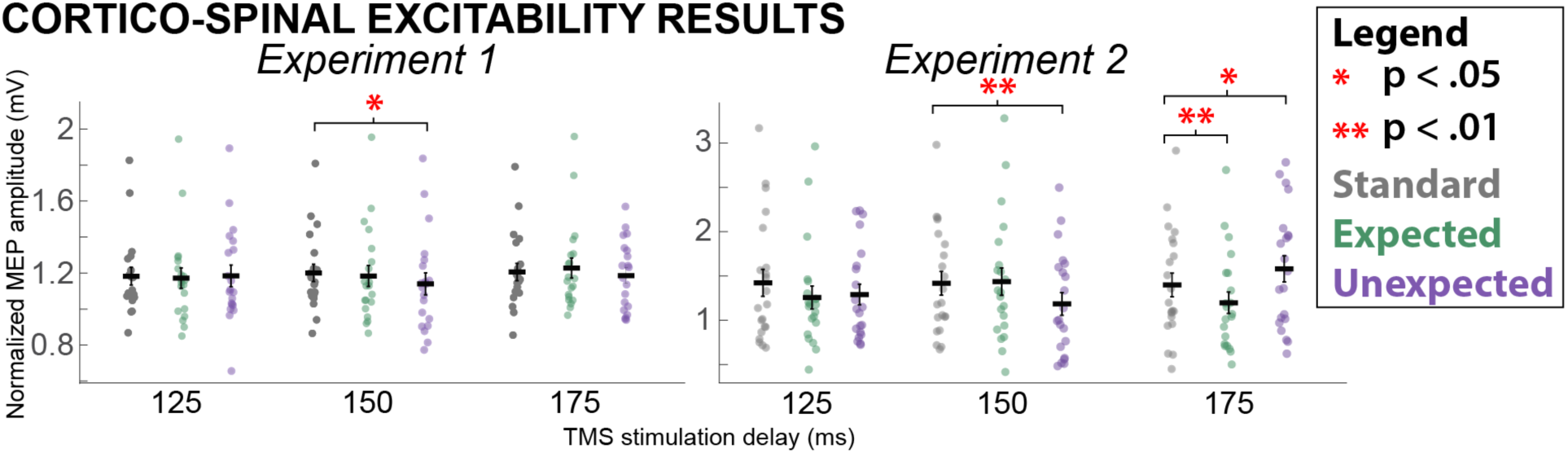
MEP results from Experiments 1 and 2, separated into trial averages (+/- standard error) by SOUND and TMS TIMING conditions. Statistically significant comparisons are noted.

In Experiment 2 (sound after imperative stimulus), no significant main effects of *SOUND* (*F*(2,20) = 2.39, *p* = .10, *η*^*2*^ = .08) or *TMS TIMING* (*F*(2,20) = 1.76, *p* = .18, *η*^*2*^ = .03) were found, but a there was a significant interaction (*F*(4,20) = 7.41, *p* < .0001, *η*^*2*^ = .37). Follow-up pairwise *t*-tests revealed that UNEXPECTED MEPs were suppressed compared to STANDARD MEPs at the 150ms delay (*t*(20) = 3.40, *p* = .003, *d* = .40), replicating our previous findings. In addition, the EXPECTED MEP was suppressed compared to the STANDARD trial MEP at the 175ms delay (*t*(20) = 2.99, *p* = .007, *d* = .39, ***Figure 2***). Moreover, at the same delay, the MEP following UNEXPECTED sounds was elevated compared to STANDARD trials (*t*(20) = 2.53, *p* = .02, *d* = .23), though unlike the above effects, this does not survive the Bonferroni-Holm correction for multiple comparisons.

## Discussion

Across two experiments, we investigated whether infrequent but expected events induce a non-selective suppression of the motor system, similarly to unexpected events and stop-signals. Using single-pulse TMS combined with EMG of task-unrelated hand muscles while participants performed forced-choice reaction time tasks with their feet, we found that infrequent expected sounds are indeed followed by a non-selective suppression of task-unrelated motor effectors. However, we found that this is only the case when a movement is currently being initiated (i.e., when the infrequent event follows the imperative stimulus that instructs a movement). In contrast, unexpected infrequent events non-selectively suppress CSE even in the absence of movement initiation (i.e., when presented before any imperative stimulus). The latter finding replicates our previous report of non-selective CSE suppression in a verbal reaction time task when unexpected sounds were presented prior to the imperative stimulus (Wessel & Aron, 2013). Notably, the timing of the non-selective suppression of CSE after unexpected infrequent events was in line with our prior work, in that it takes place at 150ms following the onset of the sound (Wessel & Aron, 2013; Dutra et al., 2018). In turn, it is notable that non-selective CSE suppression after expected infrequent events did not occur until 175ms after the event. These findings have two primary implications, which we will now discuss in turn.

First, the results suggest that there is a qualitative difference in the non-selective suppression of the motor system that takes place after infrequent events, depending on whether these events were *unexpected* or *expected*. Specifically, unexpected events induce CSE suppression even in the absence of motor initiation, which suggests a more drastic type of inhibitory control that is not evoked by expected infrequent events. Moreover, the latency difference in CSE suppression between unexpected and expected infrequent events suggests a more rapid engagement of inhibitory control when infrequent events are unexpected. In that respect, it is interesting to observe that the respective suppressive effects of expected and unexpected infrequent events do not seem to be additive. This is evident from the fact that while expected infrequent sounds produced CSE suppression at 175ms following sound onset in Experiment 2, no suppression was observed at that time point for unexpected sounds (which showed CSE suppression at 150ms instead). If the effects of surprise and infrequency were additive, unexpected sounds should have produced suppression at both 150ms (due to the unexpectedness) and at 175ms (due to the infrequency). Instead, infrequency and unexpectedness appear to independently engage the same inhibitory process, but with different intensity and latency. This supports the theory that surprise is accompanied by a unique pattern of automatically engaged inhibitory control (Wessel & Aron, 2017).

Beyond these implications for the processing of unexpected and infrequent events, the current findings also have very important implications for the study of motor inhibition in the context of the stop-signal task. As mentioned in the introduction, recent years have seen a controversial discussion regarding the exact psychological and neural mechanisms that contribute to performance in the stop-signal task. Specifically, the notion that the ability to stop an action is not solely dependent on the efficacy of the inhibitory process itself, but also depends on the initial attentional detection of the (infrequent) stop-signal and the associated triggering of the inhibitory process has been particularly prominent (Levy & Wagner, 2011; Verbruggen et al., 2014; Erika-Florence et al., 2014; Matzke et al., 2013, 2017). This notion has spurred a fundamental discussion about which parts of the neural cascade of activity following stop-signals reflect the attentional detection of an infrequent instructed signal to stop, and which reflect the motor inhibition process itself (Aron et al., 2014; Hampshire & Sharp, 2015; Swick & Chatham, 2014). In many studies that address this question, an inferential contrast is used in which brain activity following stop-signals is compared to brain activity following perceptually identical, infrequent, expected events that do not convey a ‘stopping’ instruction (Schmajuk et al., 2006; Dimoska & Johnstone, 2008; Hampshire et al., 2010; Boehler et al., 2010; Tabu et al., 2011; Dodds et al., 2011; Chatham et al., 2012; Erika-Florence et al., 2014; Bissett & Logan, 2014; Elchlepp et al., 2015; Lawrence et al., 2015; Verbruggen et al., 2010; Waller, Hazeltine, and Wessel, 2019). In other words, those studies employ a purportedly ‘non-inhibitory’ control condition that resembles the design of our current Experiment 2, where a go-signal is followed by an infrequent expected event. The current results clearly show that presenting such infrequent, expected events after go-signals lead to an automatic engagement of non-selective motor inhibition. This is in line with our other recent work, which has shown that expected infrequent events after a go-signal lead to an incidental slowing of reaction times and elicit neural activity from the same neural generator that is engaged by stop-signals (Waller et al., 2019). Taken together, this suggests that the ‘inhibition-free’ control conditions that are used in studies to isolate attentional from inhibitory processes are not, in fact, free of inhibitory activity. Hence, a subtractive contrast between stop-trials and such control conditions will likely cancel out (at least parts of) the inhibitory process, instead of isolating it. Therefore, these subtractive contrasts might operationalize other condition differences between stop-trials and control trials with infrequent signals (such as the fact that stop-trials do not include a motor response).

The current study has two shortcomings, owing to methodological limitations. First, we did not find the behavioral effects of unexpected and expected infrequent events (reaction time slowing) that are usually found in studies that use similar experimental paradigms (e.g., Dawson et al., 1982, Parmentier et al., 2008, Waller et al., 2019). This is likely due to the presence of the TMS pulses, which eliminate such behavioral effects. First, TMS of motor cortex interferes with ongoing behavior itself by interrupting the underlying motor processes (Hadipour-Niktarash et al., 2007; Cohen et al., 2009). Second, TMS pulses produce a stereotypic auditory and haptic sensation that occurs on every trial. Prior research has shown that when infrequent or surprising sounds are followed by a stereotypic, non-surprising sounds on every trial, the effect of infrequency or surprise on behavior is greatly reduced (Parmentier, 2014; Parmentier et al., 2008). This outcome is unavoidable in studies using TMS to probe the effects of unexpected events on motor excitability. A second shortcoming of the current study is we did not use an active baseline in the inter-trial interval during the task (unlike e.g., Wessel & Aron, 2013). The introduction of such baseline trials would have further elongated an already tedious and tiring task for the subjects, who had to respond to more than 800 very simple stimuli for more than 45 minutes to provide sufficient number of trials in all three conditions. Therefore, it is – strictly speaking – not possible to ascertain whether infrequent events study lead to a suppression of the MEP below a task-baseline based on the current data, or whether they merely suppress CSE compared to frequent events. However, since we know from previous study (Wessel & Aron, 2013) that unexpected infrequent events indeed suppress CSE below active baseline, one could extrapolate that the same would be true for the expected infrequent events in the current study (the CSE suppression that occurred at 175ms after expected sounds was indistinguishable in amplitude from the CSE suppression that occurred at 150 following unexpected sounds). Nevertheless, this hypothesis would necessitate independent validation.

Finally, the differential timing of CSE suppression found for unexpected and expected infrequent events provides some interesting aspects for future study. There are several potential explanations for this difference in timing. It is widely believed that non-selective CSE suppression is due to the engagement of a specific fronto-basal ganglia inhibitory pathway (Aron, 2011; Jahanshahi et al., 2015; Neubart et al., 2010; Wiecki & Frank 2013; Wessel et al., 2016; Wessel & Aron, 2017; Kelley et al., 2018; Wessel et al., 2019). It is unclear whether the timing difference between unexpected and expected infrequent events found here is due to differences in subcortical processing in the basal ganglia, or because of differences in the cortical processes that trigger the basal ganglia processes. One property of this proposed fronto-basal ganglia pathway underlying non-selective CSE suppression is its ostensible hyper-direct, mono-synaptic connection from the cortical areas that trigger the inhibitory process into the basal ganglia structures that implement the actual inhibition (Nambu et al., 2002; Parent and Hazrati, 1995; Chen et al., 2020). If this circuit is indeed as hard-wired and low-level as believed, differences in cortical processing related to the triggering of the inhibitory process are perhaps more likely to account for the differences in timing of the CSE suppression between expected and unexpected infrequent events. Indeed, classic EEG studies of such events do indicate that while unexpected infrequent events evoke a fronto-central P3a waveform, expected frequent events evoke a slower-latency, more posterior P3b (Courchesne et al., 1975; Friedman et al., 2001; Comerchero & Polich, 1999), which would suggest differences in cortical processing. Future studies could test whether both of these potentials reflect the activity of different cortical pathways that detect unexpected and expected infrequent events, respectively, but ultimately both produce inhibition mediated via the same basal ganglia circuit.

In summary, we here find that infrequent events produce a non-selective suppression of the motor system, even when they are expected. However, this suppression of the motor system is qualitatively different than the suppression observed after unexpected events (which manifests with lower latency and is also observable in the absence of motor preparation). The presence of this frequency-related inhibitory effect poses an important challenge for studies of motor inhibition that seek to produce conditions that do not include inhibitory activity. Finally, the current results show that surprise caused by unexpectedness has unique effects on the motor system that are not attributable to the relative frequency of an event alone.

## Acknowledgements

The authors would like to thank Nathan Chalkley, Kylie Dolan, and Brynne Dochtermann for their help with data collection.

## Ethics approval

University of Iowa Institutional Review Board (#201711750)

## Code/data availability

All data and code will be made publicly available on the OSF.

